# Spontaneous eye blinking is influenced by activity, synchronized with attentional breakpoints, but not modulated by social relevance in Barbary macaques

**DOI:** 10.1101/2025.08.14.670289

**Authors:** J. Ostner, R. Honnavara, C. Bruchmann, O. Schülke

**Author notes:** corresponding author: Julia Ostner, Department for Behavioral Ecology, Georg-August-Universität Göttingen, Kellnerweg 6, 37077 Göttingen, Germany.

## Abstract

Spontaneous eye blinking is a ubiquitous behavior in animals including humans necessary for lubricating the ocular surface and preventing dryness. Beyond this functional role, eye blinking also provides a window into an animal’s cognitive state and attention allocation. Here, we investigated in 13 female Barbary macaques the modulation of spontaneous eye blinking during two naturally occurring activities differing in their attentional demand (resting and allo-grooming) and additionally assess the influence of social relevance of the interaction on attention allocation. Eye blink rates were significantly lower during grooming compared to resting, suggesting increased attention during this cognitively more demanding task. Dominance rank difference and affiliative relationship strength between the groomer and groomee did not additionally influence eye blink rate. Blinking was timed to coincide with ingestion events during grooming, which may serve as explicit attentional breakpoints. By systematically timing blinks with periods of decreased visual demand, macaques effectively minimized information loss during the non-visual phase of the grooming process. Our study provides insights into the regulation of spontaneous eye blinking in nonhuman animals using a non-invasive tool for the study of visual attention and cognitive load.

## Introduction

Under normal conditions the rate of spontaneous eye blinking, i.e. involuntary and non-reflexive, is about 15-20 blinks per minute with each blink lasting 100-200 milliseconds on average despite substantial interindividual variation (Eckstein et al., 2017; Nakano et al., 2013; VanderWerf et al., 2003). Blinking functions in lubrication, thus preventing the ocular surface from drying out (Korb et al., 1994), yet it necessarily interrupts visual input (Bristow et al., 2005). As the average blink rate exceeds the rate necessary for efficient lubrication of the cornea by far, the blink rate can be modulated in response to the cognitive demand of a task (Cornelis et al., 2025; Ponder & Kennedy, 1927). Indeed, the rate of spontaneous blinks is reduced both in within- and between subject experimental designs during more cognitive demanding tasks (Ichikawa & Ohira, 2004; Oh et al., 2012; Siegle et al., 2008), tasks requiring focused visual attention, such as tracking (De Jong & Merckelbach, 1990), during reading (Abusharha, 2017; Bentivoglio et al., 1997), or when watching relevant, memorable scenes in a movie (Shin et al., 2015). Blink rate inhibition has also been demonstrated in more complex real-life situations such as driving a car and aircraft control (Ahlstrom & Friedman-Berg, 2006; Faure et al., 2016), indicating cognitive regulation of the rate of spontaneous blinks depending on attentional demand.

Modulation of the rate of spontaneous eye blinking depending on the task at hand has also been shown in nonhuman animals. As blinking interrupts visual input, suppression of blinks occurs in individuals exposed to danger requiring increased attention. Wild crows reduce blinking to fix their gaze upon a possible danger, i.e. a threatening human experimenter (Cross et al., 2013), and wild anubis baboons (Matsumoto-Oda et al., 2018) and red deer (Rowe et al., 2023) have lower eye blink rate in smaller groups with a presumably higher predation pressure and increased need to be vigilant. Blink rate is also lower in younger more peripheral individual baboons compared to older more central, presumably safer, group members (Matsumoto-Oda et al., 2018). Reflecting findings in human social attention and the suppression of blinking during cognitively demanding, or more relevant tasks, great-tailed grackles strategically inhibit their blinking behaviour during flight compared to before and after (Yorzinski, 2020a) and Japanese macaques reduce blinking during allo-grooming compared to resting periods (Hikida, 2022). Rhesus macaques decrease blinking when watching videos of conspecifics compared to a control situation (blank screen) and the reduction was further amplified by visual relevance (more than one monkey, more complex environment on screen; (Ballesta et al., 2016)). The modulation of spontaneous blink rate as a function of visual demand and relevance in nonhuman animals, thus, is consistent with findings in humans showing an inverse relationship between eye blink rate and attention (see above).

To minimize the loss of visual information during blinking and to allow attentional disengagement, blinks can be strategically timed to coincide with attentional break points (Nakano et al., 2009, 2013). Indeed, blinking tends to occur at punctuations during reading (Cornelis et al., 2025), at a pause by the speaker while listening, or the conclusion of an action and the repeated presentation of a similar scene while watching videos (Andreu-Sánchez et al., 2021; Nakano et al., 2009). Again replicating findings in humans, nonhuman animals align their blinks to break points: grackles are more likely to blink during impact compared to in-flight, potentially to compensate for the low blink rate prior to impact (Yorzinski, 2020a), peacocks time their blinks with large gaze-shifts (Yorzinski, 2016), and grooming Japanese macaques synchronize blinks with the ingestion of ectoparasites, i.e. the period during allo- grooming requiring the least visual attention (Hikida, 2022). These data collectively indicate a regulation of blinking in such a way that as little information as possible is lost while at the same time the necessary biological function of lubrification is maintained.

Continuous visual monitoring is required for situations that demand immediate action because they pose a threat, such as exposure to a predator or other potential dangers, yet also for tasks that need close visual attention, such as grooming. Allo-grooming is a widespread element in the behavioral repertoire of social mammals serving both hygienic and social functions (Grueter et al., 2013). It is defined as the repeated stroking through an individual’s fur thereby locating and removing ectoparasites, their eggs, and other particles, interrupted by ingestion of these particles (Jaeggi et al., 2017). Given the small size of parasites and other particles, efficient allo-grooming requires a high level of visual attention with ingestion events serving as potential attentional break-points (Hikida, 2022).

In addition to parasite removal, grooming serves pivotal social functions. It is the main affiliative behavioral pattern in nonhuman primates, distributed selectively among partners, and instrumental for the formation and maintenance of social bonds (Henzi et al., 1997; Silk, 2002) increasing cooperation and support among grooming partners (Henzi & Barrett, 1999; Schino, 2007). Therefore, the value of available grooming partners within a social group varies depending on the affiliative relationship a groomer has with their potential groomee; the closer the relationship, the more supportive it is and thus the higher the relationship value. On another dimension, individuals vary in their social status, i.e. dominance rank, where a higher-ranking individual is better able to provide access to resources and is a more powerful ally in an agonistic conflict (Seyfarth, 1977). Consequently, grooming activities gain additional relevance based on the affiliative relationship and dominance status of partners, which may moderate the degree of investment and thus visual attention given by the groomer to the task at hand.

In this study, we used the rate of spontaneous eye blinks during natural behaviors to quantify visual attention to grooming interactions and the potential moderating role of relational aspects in Barbary macaques. We specifically predicted the eye blink rate to be decreased during grooming compared to a resting control and this effect to increase in size with increasing affiliative relationship strength and increasing rank difference between the groomer and the groomee. Additionally, we predicted, that in order to minimize the loss of visual information due to blinking, individuals will strategically time their blinking to coincide with ingestion events.

## Methods

### Study site and subjects

Study subjects belonged to a multimale-multifemale group of Barbary macaques living in semi-free ranging conditions at Affenberg Salem, a 20ha forested enclosure in Southern Germany (https://www.affenberg-salem.de/en/). Animals fed on naturally available vegetation and insects, and were additionally provisioned once a day with fruit, vegetables, and grains, and had *ad libitum* access to water (de Turkheim & Merz, 1984). Subjects for this study were all 13 adult (> 5 years) female Barbary macaques of group H (total group size: 41 individuals including 14 adult males, 14 immatures) and observations took place from June 2024 to August 2024. All focal females were reliably recognizable by natural facial and body features, genital swellings, and inner leg tattoos. Data on individual age were available from Affenberg Salem records.

### Data collection

Eyeblink data were collected during focal animal sampling spread evenly across all hours of the day (Altmann, 1974). Whenever the focal animal started resting (defined as being stationary, not feeding, sleeping, or engaged in a social interaction) or actively allo-grooming another adult female (defined as brushing through another’s individual’s fur and removing ectoparasites, dirt, or dead skin (Jaeggi et al., 2017)), we video-recorded the focal female’s face using a Sony CX240E Handycam. The recording stopped when the focal female displayed another behavior for > 15 sec, closed her eyes, or moved the face preventing an appropriate camera angle. In addition, to eyeblink occurrences, we scored ingestion events (defined as the groomer picking particles (dirt, skin, ectoparasites) from the fur of the groomee and immediately ingesting those).

In addition to focal animal sampling, we conducted group scan and ad libitum sampling for the quantification of affiliative relationship strength and dominance rank (Altmann 1974). Group scans were conducted simultaneously by 2-3 observers 4-6 times per day with at least one hour between consecutive scans (total number of group scans = 430). A scan contained only adult individuals, lasted maximally 12 minutes, and included on average 23.3 of the 27 adult individuals (range = 17 – 27). The subject’s activity (grooming, resting, feeding, travelling) as well as partner identity in social interactions (grooming, resting in body contact, 1.5m proximity) were recorded. Throughout the study period, we recorded ad libitum data on decided dyadic agonistic conflicts between two adult females, i.e. conflicts with a clear loser showing submission, either unprovoked or following an aggression by the opponent (total 992 decided conflicts).

### Behavioral data analysis

Eye blink data were extracted from focal animal protocols using the video logging software BORIS version 8.16.13 (Friard & Gamba, 2016). Two state (grooming, resting) and two point (eye blink, ingestion) behaviors were coded. Grooming and resting states shorter than 30sec were excluded from analysis. Across the thirteen focal females, a total of 151 resting recordings with a total duration of 240.78 min (mean duration = 1.59min, range= 0.5 – 7.08) and 229 allo-grooming recordings with a summed duration of 330.99 min (mean duration = 1.45min, range= 0.5 – 4.75) were coded.

A dominance hierarchy was established based on the outcome of 992 dyadic decided conflicts based on the Elo Rating method (Albers & De Vries, 2001) using the package EloRating (v0.46.11 (Neumann et al., 2011)) using the default settings. Elo score at the end of the observation period was used as a measure of dominance success. Affiliative relationship strength was quantified as the Dyadic Composite Sociality Index (DSI, (Silk et al., 2006, 2013)) from the group scan data set (N = 430, (McFarland et al., 2017)) using counts of two positively correlated affiliative behaviors, allo-grooming and being in close (1.5m) spatial proximity. The correlated metrics were integrated into the DSI (Silk et al., 2013) as follows: 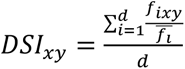.The DSI was composed of the rate of a behavior i of a dyad xy, divided by the mean rate of this behavior i of all dyads. The variable d represents the number of behaviors included, here two. Using the group scan data, matrices of dyadic grooming and proximity interaction frequencies were calculated and divided by the number of scans in which both members of a dyad were observed in the same scan. In this way, the adjusted frequency of the behavior (i) is obtained, which controls for biases due to different observation frequencies, thus ensuring normalization of the data (Young et al., 2017). The DSI by definition has a mean of one and increasingly high values indicate that dyads had been increasingly more often in affiliative contact for increasingly more time than the average dyad in the group, while a dyad with low values spends less than average time affiliating.

### Statistical analyses

#### Predictors of eye blink rate

To estimate the extent to which eye blink rate differed between grooming and resting (model 1), we fitted a Generalized Linear Mixed Model (GLMM) (Baayen et al., 2008). Into this model we included a fixed effect of activity type and also one for Elo score of the groomer to control for its effect. We further included a random intercepts effect for the ID of the groomer. To avoid an ‘overconfident’ model and keep type I error rate at the nominal level of 0.05 we included a random slope of activity type (Barr et al., 2013; Schielzeth & Forstmeier, 2009). Finally, to control for the duration of the observation, we included an offset term for observation duration (log-transformed, base e; (McCullagh & Nelder, 1996)). Note that such a count model with observation time included as an offset term, effectively models eye blink rate (i.e., the number blinks per second). We fitted the model with a zero truncated negative binomial error distribution, since the response did not comprise a single zero, and a Poisson model was clearly over-dispersed. To avoid convergence problems, we had to refrain from estimating the correlation between the random intercept and slope. Prior to including the random slope of activity type we manually dummy coded and then centered it.

To estimate the extent to which eye blink rate correlated with the rank difference between the groomer and the groomee and with their dyadic affiliative relationship we fitted a corresponding model (model 2) this time only including observations during which grooming took place. We again fitted the model with a zero-truncated negative binomial error distribution and also included an offset for the duration of the event. As fixed effects we included Elo score difference and dyadic relationship strength between groomer and groomee. We included random intercepts effects for the ID of the groomer, the groomee, and the groomer-groomee dyad to account for individualized relations. We included random slopes of both rank difference and dyadic relationship within groomer ID and groomee ID. In order to avoid ‘cryptic multiple testing’ (Forstmeier & Schielzeth, 2011), we compared this full model with a null model which lacked dominance rank difference and dyadic relationship in the fixed effects part.

We fitted the models in R (version 4.4.3, (R Core Team, 2024)) using the function glmmTMB of the equally names package (version 1.1.10; (Brooks et al., 2017)). Prior to fitting the models, we z-transformed the covariates rank, rank difference, and dyadic relationship to a mean of zero and a standard deviation of one to ease model convergence. We estimated 95% confidence limits of model estimates and fitted values by means of parametric bootstraps (N=1000 bootstraps; function simulate of the package glmmTMB). We tested the significance of individual fixed effects predictors by dropping them one a time and comparing the likelihoods of the resulting model with that of the respective full model using a likelihood ratio test (R function drop1; (Dobson, 2002)). The full-null model comparison also utilized a likelihood ratio test. In order to determine model stability, we dropped each individual level of each individual random effects factor, fitted the full model each of the subsets, and finally compared the range of estimates obtained with those obtained for the respective model (1 or 2) on the full data set. This revealed both models to be of good to excellent stability. The sample analysed with model 1 comprised a total of 380 observations (229 grooming, 151 resting events) of 13 focal females, and the sample analysed with model 2 comprised the subset of 229 grooming observations including 56 dyads. In neither of the two models the response was over dispersed given the model (dispersion parameters, model 1: 0.82; model 2: 0.92).

### Co-occurrence of eye blinks and ingestion events

To test whether the occurrence of eye blinks and ingestion events was temporally associated, we used a permutation test (Adams & Anthony, 1996; Manly, 2007) which we applied to each observation period separately. To this end, we first determined for each ingestion event the absolute time lag to the nearest eye blink, then averaged this across the eye blinks of the given observation period, and took this as the test statistic. We then permuted the eye blink events by shuffling the intervals between them. That is, the moments of the first and last eye blinks, the number of eye blinks, and the distribution of time lags between them remained unaltered. We conducted 1000 permutations whereby we included the original data as one permutation, and each time determined the test statistic. We finally determined the one tailed p-value as the proportion of permutations revealing a test statistic at least as small as the original data. Since we applied such a test for 196 observation periods, an error level correction for multiple testing was needed. We applied an informal correction, resting on the fact that a number of P-values obtained from processes for which the null hypothesis is true are uniformly distributed in the interval from zero to one. Hence, plotting the cumulative relative frequency distribution of P-values against the P-values should reveal a more or less straight line from x=0 and y=0 to x=1 and y=1. In case of a temporal association of eye blinks with ingestions, the P-values will concentrate at lower values, and their cumulative relative frequency distribution will increase faster than expected at smaller P-values.

## Results

### Predictors of eye blink rate

The type of activity a female was engaged in influenced her eye blink rate, as blink rate during active grooming (mean ± STD = 0.18 ± 0.07 blinks per second, N = 229) was clearly lower than during resting (mean ± STD = 0.29 ± 0.12, N = 151). The control predictor (groomer’s dominance rank) was not significant (model 1; Table 1; Figure 1).

**Figure 1:**
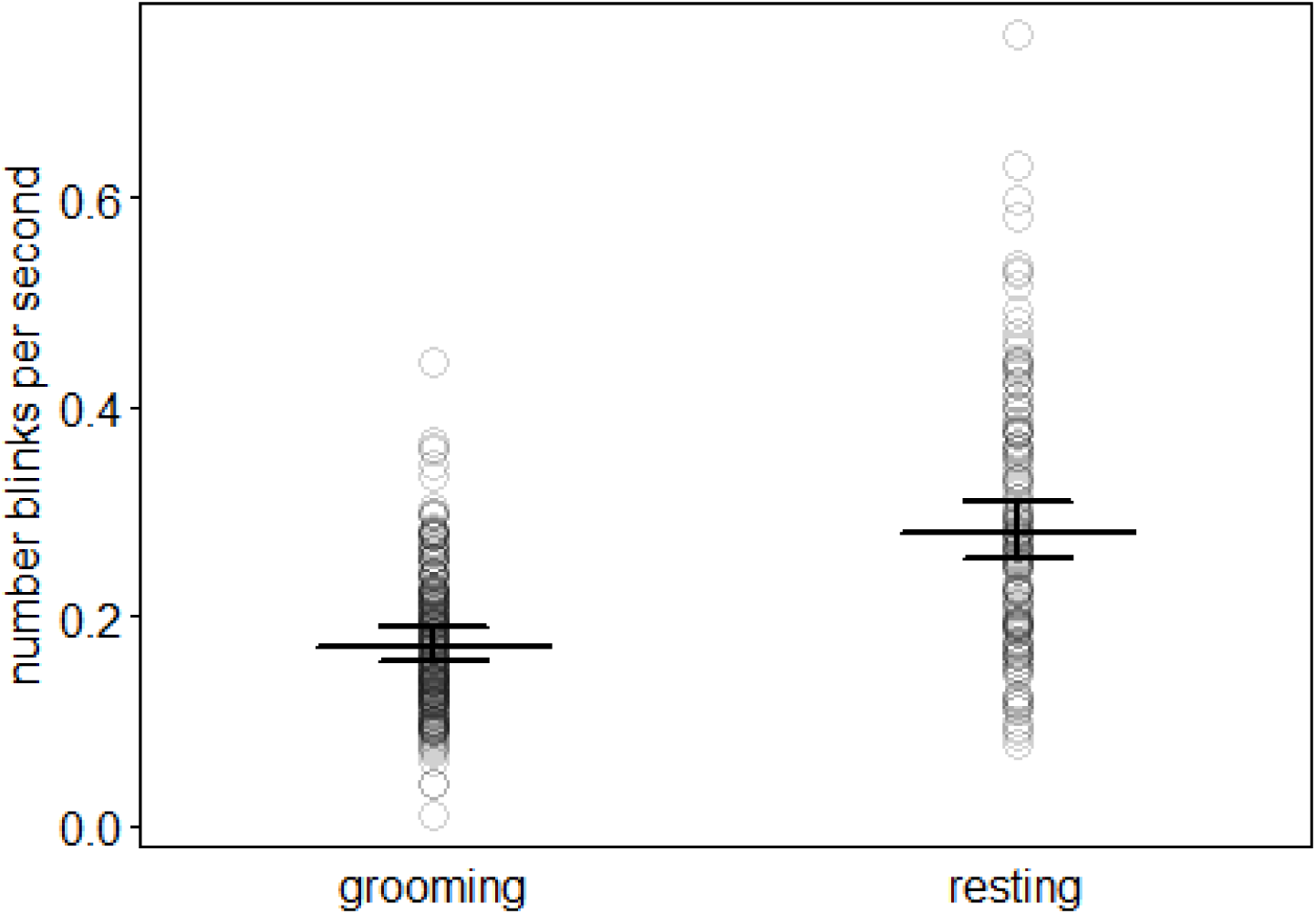
Number of eye blinks per second during the two activity types, grooming and resting, respectively. Dots show individual observations and horizontal line segments with error bars depict the fitted model (model 1) and its 95% confidence limits for an individual with an average Elo score (dominance rank).

**Table 1:** Effect of activity type on eye blink rate (GLMM, model 1)

| Term | Estimate | SE | p-value | 95% CI | ChiSq | Min | Max |
| --- | --- | --- | --- | --- | --- | --- | --- |
| **Intercept** | -1.76 | 0.05 |  | [-1.86, -1.67] |  | -1.78 | -1.72 |
| **Activity type (resting)<sup>1</sup>** | 0.49 | 0.04 | <0.001 | [0.42, 0.57] | 36.91 | 0.47 | 0.51 |
| **Subject rank<sup>2</sup>** | -0.05 | 0.04 | 0.18 | [-0.13, 0.03] | 1.47 | -0.13 | -0.02 |

*Note:* Indicated are model estimates, together with their standard errors, significance test, 95% confidence limits, likelihood ratio test, and the range of estimates obtained when excluding individuals one at a time.
<sup>1</sup>Activity type was dummy coded with grooming being the reference level
<sup>2</sup>Subject rank was z-transformed to mean = 0 and SD = 1; mean and SD of original rank were 610.97 and 326.57, respectively

In case of model 2, the full-null model comparison did not reveal significance (χ^2^ = 0.485, df = 2, P = 0.785). Correspondingly, neither dyadic affiliative relationship strength nor dyadic dominance rank difference were significant predictors of eye blink rate during active allo- grooming (Table 2; Figure 2).

**Figure 2:**
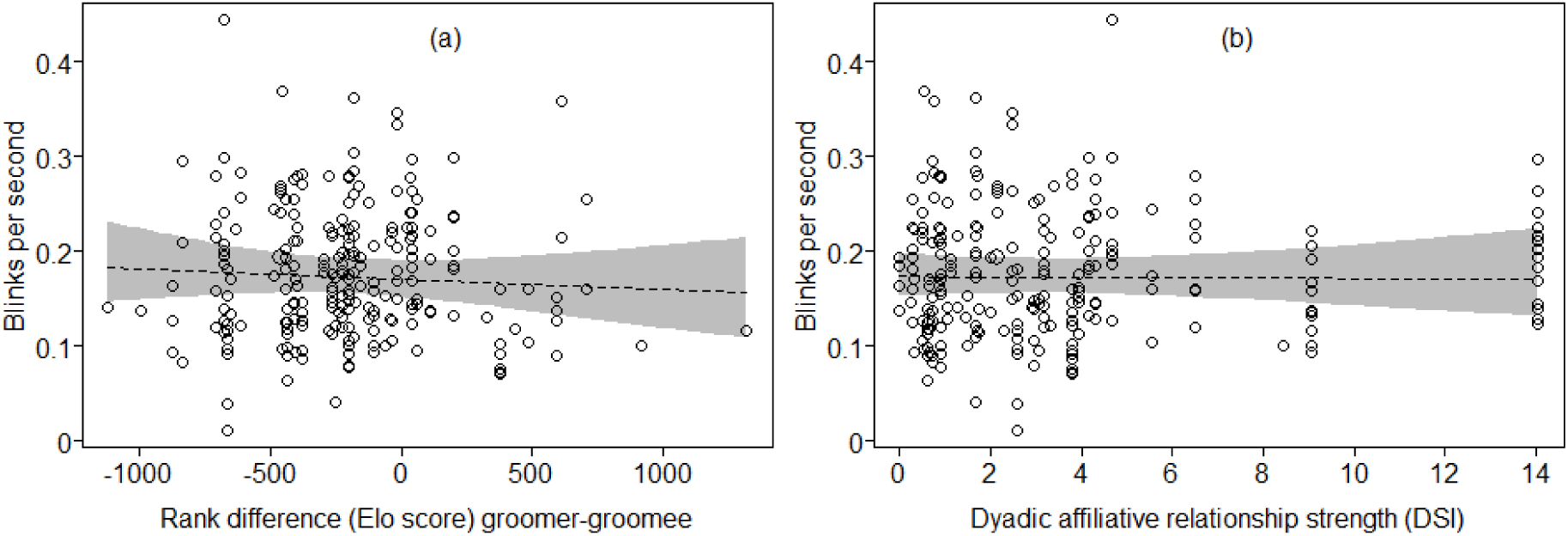
Number of eye blinks per second as a function of (a) dominance rank (Elo score) difference and (b) dyadic affiliative relationship strength (DSI) between groomer and groomee. Dots show individual observations and the dashed line with the grey area depicts the fitted model (model 2) and its 95% confidence limits for a dyad with (a) an average relationship strength and (b) an average rank difference respectively.

**Table 2:** Effect of dominance rank difference (Elo score) and dyadic affiliative relationship strength (DSI) on eye blink rate during grooming interactions (GLMM, model 2)

| Term | Estimate | SE | p-value | 95% CI | ChiSq | Min | Max |
| --- | --- | --- | --- | --- | --- | --- | --- |
| **Intercept** | -1.76 | 0.05 |  | [-1.86, -1.65] |  | -1.78 | -1.67 |
| **Rank difference<sup>1</sup>** | -0.02 | 0.04 | 0.53 | [-0.1, 0.06] | 0.394 | -0.06 | -0.003 |
| **Relationship strength<sup>2</sup>** | -0.01 | 0.04 | 0.91 | [-0.09, 0.07] | 0.01 | -0.03 | 0.006 |

*Note:* Indicated are model estimates, standard errors, 95% confidence limits, significance tests, and the range of estimates obtained when excluding groomers, groomees, and dyads one at a time.
<sup>1</sup>Rank difference was z-transformed to mean = 0 and SD = 1; mean and SD of original rank difference were -218 and 360
<sup>2</sup>Relationship strength was z-transformed to mean = 0 and SD = 1; mean and SD of original affiliative relationship strength were 3.53 and 3.57.

### Co-occurrence of eye blinks with ingestion events

As predicted, during grooming interactions there was clear temporal association of eye blinks with ingestion: the distribution of the actually observed P-values concentrated at lower values, and their cumulative relative frequency distribution increased much faster than expected at smaller P-values (Figure 3). In fact, about 40% of the absolute time lags between an ingestion event and the closest eye blink was ≤ 0.25 seconds whereas in the randomized data less than 10% of time lags were this short (Figure 4).

**Figure 3:**
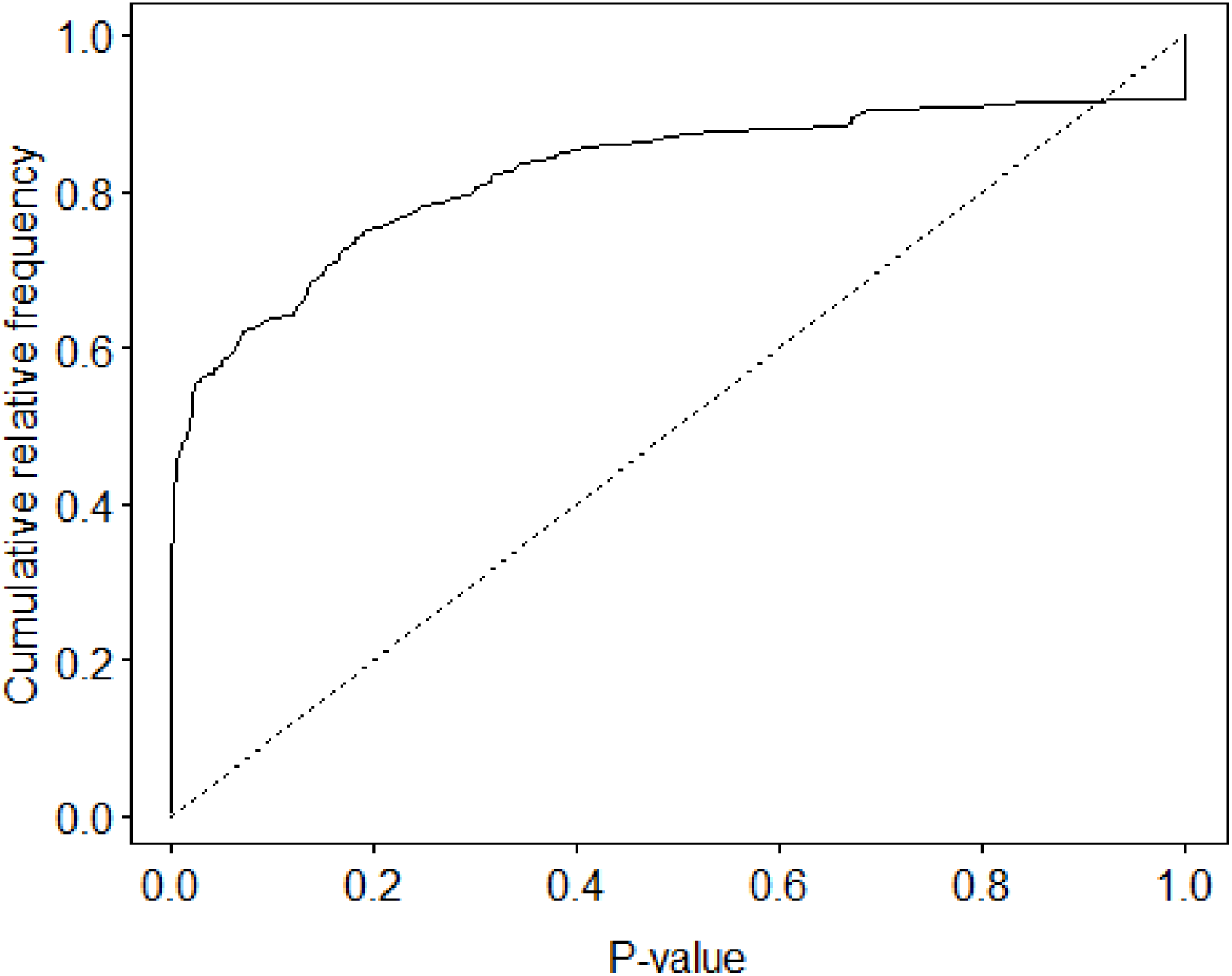
Relative cumulative frequency distribution of P-values of the temporal association of eye blinks with ingestions (solid line) and the respective expected distribution under the assumption of the true null hypothesis. Note that the distribution of the actually observed P-values was clearly biased towards smaller P-values.

**Figure 4:**
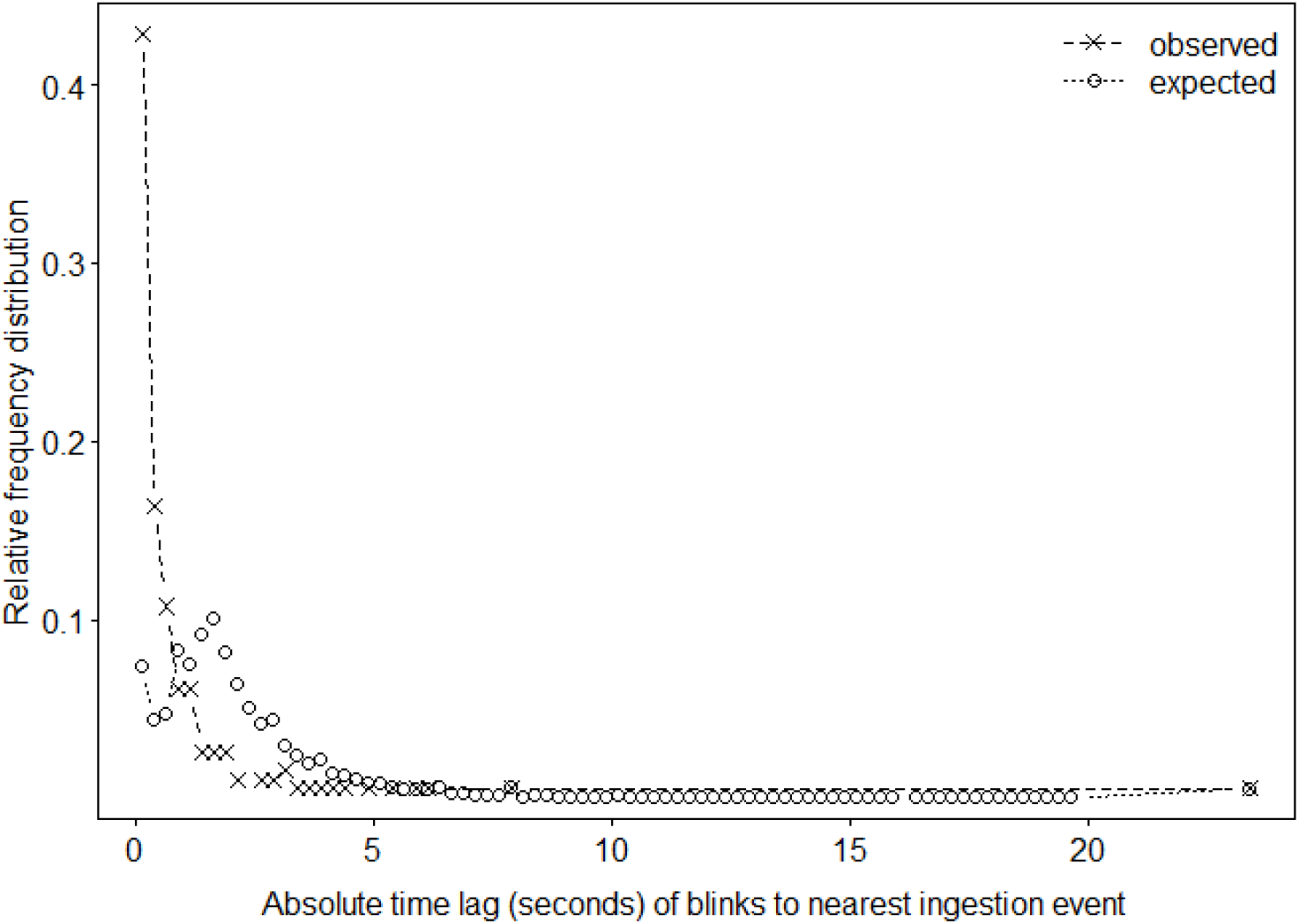
Relative frequency distribution of absolute time lags between ingestions and the closest eye blink in observed (dashed line) and shuffled (expected) data (dotted line).

## Discussion

Our study provides further evidence for spontaneous eye blink rate as an indicator of increased attentional focus in nonhuman animals consistent with findings in humans (Bentivoglio et al., 1997; Hikida, 2022; Yorzinski, 2020a)^13,20,21^. The average baseline (resting) blink rate of Barbary macaques of this study (17.4 blinks per minute) closely resembles the rate reported for the species and falls within the macaque genus range (19.5 blinks per minute for Barbary macaques, range across 11 macaque species 5.1 – 22.6 blinks per minute; (Tada et al., 2013)). We observed an inhibition of spontaneous eye blinking, and thus increased attention allocation, during grooming compared to resting, potentially to increase information absorption during the attention-demanding task of perceiving and picking up very small particles (lice, eggs, dirt) in the dense fur of a conspecific. This task-related modulation of blink rates mirrors the increasing number of findings of an inverse relationship between attentional requirements and spontaneous blink rate in a range of vertebrate species: A reduced blink rate was associated with increased (perceived or real) environmental risk in deer, baboons, and several species of birds (Beauchamp, 2001; Matsumoto-Oda et al., 2018; Rowe et al., 2023; Yorzinski, 2020a; Yorzinski et al., 2021), with visual attention to increasingly complex social stimuli in rhesus macaques (Ballesta et al., 2016) and as in our study with a visually demanding naturally occurring social task (allo-grooming) in Japanese macaques (Hikida, 2022).

Contrary to our expectation, eye blink rate inhibition during grooming was not modulated additionally by social factors. Grooming in cercopithecine primates is often directed up the hierarchy in exchange for access to rank-related benefits such as support in conflicts or tolerance for access to food (Henzi & Barrett, 1999; Schino, 2007; Seyfarth, 1977). Given the linear dominance hierarchy among the adult females of this study, we predicted increased attentional focus with an increasing dominance rank differential between the groomer and groomee, a prediction that was not supported by our data. While we can only speculate why rank difference did not additionally modulate blink rate inhibition, we propose three potential, non-exclusive explanations for the lack of an effect of rank difference. First, while grooming undoubtedly requires focused visual attention explaining the overall reduction in eye blinking during this activity compared to resting, an additional modulation may not be needed if there is already a ceiling effect of performance improvement. Missing visual input momentarily may not be as detrimental as for example during threat detection (Schülke et al., 2020; Yorzinski et al., 2021) or when perceiving facial signals of conspecifics (Ballesta et al., 2016). Second, grooming someone far higher in rank may presents a highly vulnerable and tense social situation for the groomer. Anxiety and tension has been shown to increase eye blink rate both in humans (Giannakakis et al., 2017) and nonhuman animals (Ballesta et al., 2016; Hikida, 2022; for an opposite effect in horses see Merkies et al., 2019), maybe due to overriding attentional needs by socioemotional factors (Ballesta et al., 2016). These contrasting effects of increased attention due to partner value and increased need for attentional disengagement may then cancel each other out. Third, the increasingly high value of a partner ranking high above a groomer may have been overestimated in this food provisioned setting with *ad libitum* access to high quality resources available to all individuals regardless of their relative rank position (de Turkheim & Merz, 1984) and the generally relaxed dominance style of the study species (Thierry, 2007, 2022), reducing the degree of rank-related disparities and consequently the need to barter with a high ranker. Consistent with this, individual dominance rank both during resting and grooming did not predict eye blink rate (also not in a similar study in Japanese macaques, Hikida, 2022)). Future studies are needed to conclusively address the role of rank asymmetry in social interactions for attention allocation.

In addition to the rank difference between the grooming partners, we also investigated the effect of their affiliative relationship strength, expecting a stronger attentional focus with increasing strength. As with rank difference this social predictor had no effect on the eye blink rate during grooming. The same argument as above, a ceiling effect, no need for additional attention on grooming performance, holds here as well. In addition, our data show considerable within-dyad variation in eye blink rate (Figure 2b) which hints at relevant yet unmeasured factors driving this variation. Climatic factors such as wind and rain are known to increase eyeblink rate in birds and humans (Nakamori et al., 1997; Yorzinski, 2020b; Yorzinski & Argubright, 2019). Similarly, blinking occurs more often during head movements in primates and birds (Beauchamp, 2001; Gandhi, 2012; Yorzinski, 2016) and also with increasing tasks duration and associated mental fatigue (Maffei & Angrilli, 2019). While we did not record eye blink data during rain or heavy winds and, thus, these climatic factors should not have affected our results, other variables such as head movements and tasks duration were not measured and consequently not included into our model, may have possibly masked an effect of dyadic relationship strength on eye blinking in our study.

Blinking was temporally synchronized with ingestion events, that may serve as explicit attentional breakpoints during grooming (Nakano et al., 2013). When mouthing debris, lice, or lice eggs collected from the groomee’s fur, visual attention can be momentarily released to then be re- focused when the visual task resumes. By systemically timing blinks with periods of decreased visual demand, information loss is minimized. Similar patterns of attentional disengagement timed with implicit or explicit attentional breakpoints have been found in humans (Nakano et al., 2013) and also in non-human animals, where blinking is synchronized with head movements during feeding in chickens (Beauchamp, 2001), gaze shifts in peacocks (Yorzinski, 2016), and most closely resembling our study with ingestion events during grooming in Japanese macaques (Hikida, 2022). By systematically modulating the timing of blinking to coincide with a period of lower cognitive load, female macaques of our study effectively minimized information loss during a non-visual phase of the grooming process while by renewal of the tear film also increasing the acuity of vision for the next visually more demanding period when blinking is suppressed (Ang & Maus, 2020).

Our study adds to the still limited body of research on patterns of spontaneous eye blinking as a measure of visual attention and cognitive load in nonhuman animals. Consistent with other studies, Barbary macaques suppressed eye blinking during demanding tasks and aligned blinks with attentional breakpoints, thereby maximizing information gain. While we extend previous work by investigating the potentially modulating effects of relational factors (dominance and affiliative relationships), our study also has clear limitations. Additional factors known to influence eye blink pattern, such as head movements and gaze shifts (Beauchamp, 2001; Yorzinski, 2016) or task duration (Maffei & Angrilli, 2019), should be included in future studies on visual attention during natural occurring interactions.

## Data availability

Data are publicly available at GWDG Data repository GRO.data: https://doi.org/10.25625/HMPWNJ.

## Acknowledgements

We thank Roland Hilgartner for permission to work at Affenberg Salem, Mamisolo Hilgartner and the staff of Affenberg Salem for the help provided during data collection and Roger Mundry for statistical advice.

## Ethics Statement

The study was purely observational, strictly non-contact/non- invasive, and adhered to the guidelines for the treatment of animals in teaching and research of the ASAB Ethical Committee/ABS Animal Care Committee SAB (2023) and the EU directive 2010/63/EU.

